# Improving metazoan biodiversity inventories associated with rocky subtidal habitats of the North Colombian Pacific through eDNA metabarcoding and DNA barcodes

**DOI:** 10.64898/2026.08.06.743172

**Authors:** Vanessa Yepes-Narváez, Alejandro Rodríguez-Sánchez, Mayra Atencia-Galindo

**Author notes:** Corresponding author: (VYN). All authors contributed equally to this work.

## Abstract

The marine biodiversity inhabiting rocky shores in the Colombian Pacific remains largely undocumented, primarily due to geographic isolation, logistical challenges, and socio-political constraints. To address the existing knowledge gap, we conducted an expedition to enhance baseline biodiversity knowledge in rocky shores by integrating multiple complementary approaches, including visual censuses, specimen collection with morphological identification, environmental DNA (eDNA) metabarcoding and DNA barcodes. eDNA samples were collected at four coastal sites adjacent to rocky substrates, along with biological specimens obtained from fourteen locations through SCUBA diving at depths ranging from 1 to 25 meters. Tissue samples were subjected to genomic DNA isolation, followed by the generation and validation of cytochrome c oxidase subunit I (COI) barcode sequences, which were subsequently corroborated through taxonomic assessment to ensure accurate species identification. eDNA metabarcoding analyses yielded over 7 million high-quality sequence reads. Although taxonomic resolution at the species level was constrained by the limited completeness of reference sequence databases, a total of 106 species and 83 families were successfully identified, predominantly within the classes Actinopteri, Chondrichthyes, and marine mammals. From the 769 specimens obtained we generated 871 sequences, including 414 validated COI barcodes representing 76 species across 64 families. The integration of DNA barcoding and eDNA approaches resulted in over 1,400 taxonomic detections spanning five phyla, with only six species shared between methodologies. Richness and diversity varied among sites, and revealed significant differences along the coastline between Jurado and Cupica Gulf. All sequences were deposited in BOLDsystems database under the CCBIO project and were visualized through OBIS and GBIF databases. These findings provide the first molecular-based baseline for rocky shore biodiversity in the Colombian Pacific, highlighting the value of integrative approaches for monitoring and conservation.

## Introduction

Rocky shore ecosystems in the Eastern Tropical Pacific are structurally complex habitats composed of hard geological or biological substrates that sustain diverse assemblages of marine fauna and flora [1–3]. Their heterogeneity in slope, substrate type, and microhabitat availability promotes high species richness and intricate trophic interactions [4,5,2]. These ecosystems are divided into intertidal and subtidal zones, each with distinct environmental characteristics and communities. Subtidal zones in particular, maintain stable conditions and continuous resource availability, supporting greater structural complexity and biodiversity, whereas intertidal zones experience extreme environmental fluctuations that impose strong selective pressures shaping species distribution and composition [6–8].

In Colombia, marine biodiversity is recognized as a national heritage of high ecological, cultural, and economic significance, forming a cornerstone of environmental policy and sustainable development [9]. Although marine ecosystems encompass nearly half of the national territory, research in this field remains geographically uneven, with a historical concentration in the Caribbean region. In contrast, the northern Pacific coast, has received minimal scientific attention due to its remoteness, limited infrastructure, and socio-political constraints, resulting in a severe lack of baseline data on species composition, distribution, and ecological dynamics [10–11].

The northern Colombian Pacific is part of the Choco biogeographic region, a recognized biodiversity hotspot, ranked among the 25 most important global centers of endemism [12–13]. While terrestrial biodiversity has been well documented for decades, marine biodiversity remains poorly studied. Despite rocky shores cover approximately 636 km of the Colombian Pacific coastline [14], research on their subtidal habitats remains limited compared to other ecosystems such as coral reefs and mangroves [15]. Existing studies, concentrated mostly in southern Pacific areas such as, Gorgona Island, Malaga bay and Malpelo Flora and Fauna Sanctuary [16–18]), and have focused on taxonomic inventories, leaving major gaps in knowledge on species distribution, ecological interactions, and functional roles [19,20,6,21]. This lack of baseline data constrains our ability to assess biodiversity responses to environmental change and anthropogenic pressures, hindering effective management strategies.

Recent surveys in northern Pacific (Choco department) indicate high biodiversity associated with rocky reefs, including sponges [22], echinoderms [23], macroalgae [24], and mobile invertebrates [25], with numerous new records for Colombia, highlighting the potential for undiscovered diversity and the urgent need for systematic research to inform conservation and sustainable management. The Jurado-Cupica in the northermost Colombian Pacific region exemplifies this gap, as marine ecosystems and species records remain poorly characterized and incomplete with only limited records from Punta Cruces and Cabo Marzo [26]. This gap reflects sampling biases and an unfinished baseline of marine biodiversity.

To address these limitations, we applied DNA barcoding and environmental DNA (eDNA) as powerful approaches to complement taxa richness and composition estimations contributing to biodiversity assessment in the coastline between Jurado district and Cupica Gulf. These techniques enabled a larger species identification from minimal biological tissues and genetic traces in seawater samples, facilitating research in such remote and logistically challenging area. Results allowed a more comprehensive species inventories, detection of cryptic taxa, and proposed a cost-eficient tool for ecosystem monitoring, as well as the generation of genetic datasets to inform conservation management. Integrating these methodologies into marine research was therefore essential to bridge existing knowledge gaps on metazoan composition in subtidal rocky habitats and contributed to strengthen Colombia’s capacity for future sustainable management and conservation planning of its marine resources.

## Materials and methods

### Study area

The study was conducted along the 88.72km coastline between the municipality of Juradó in Punta Melo (6°57’19.031”N, 77°40’36.25”W) and the subtidal rocky cliffs of Chicocora in Cupica Gulf (6°40’36.12”N, 77°25’51.69”W) in the northern Colombian Pacific (Chocó Department) (Fig 1). The area features a heterogeneous coastal geomorphology with cliffs, beaches, estuaries, and insular outcrops [27]. Rocky shore is composed of escarpments of sedimentary and volcanic rocks alternated with unconsolidated deposits. Oceanographic dynamics driven by a mixed tidal regime and high rainfall, intensify erosion and shape complex intertidal habitats [28, 26, 29].

**Fig 1.**
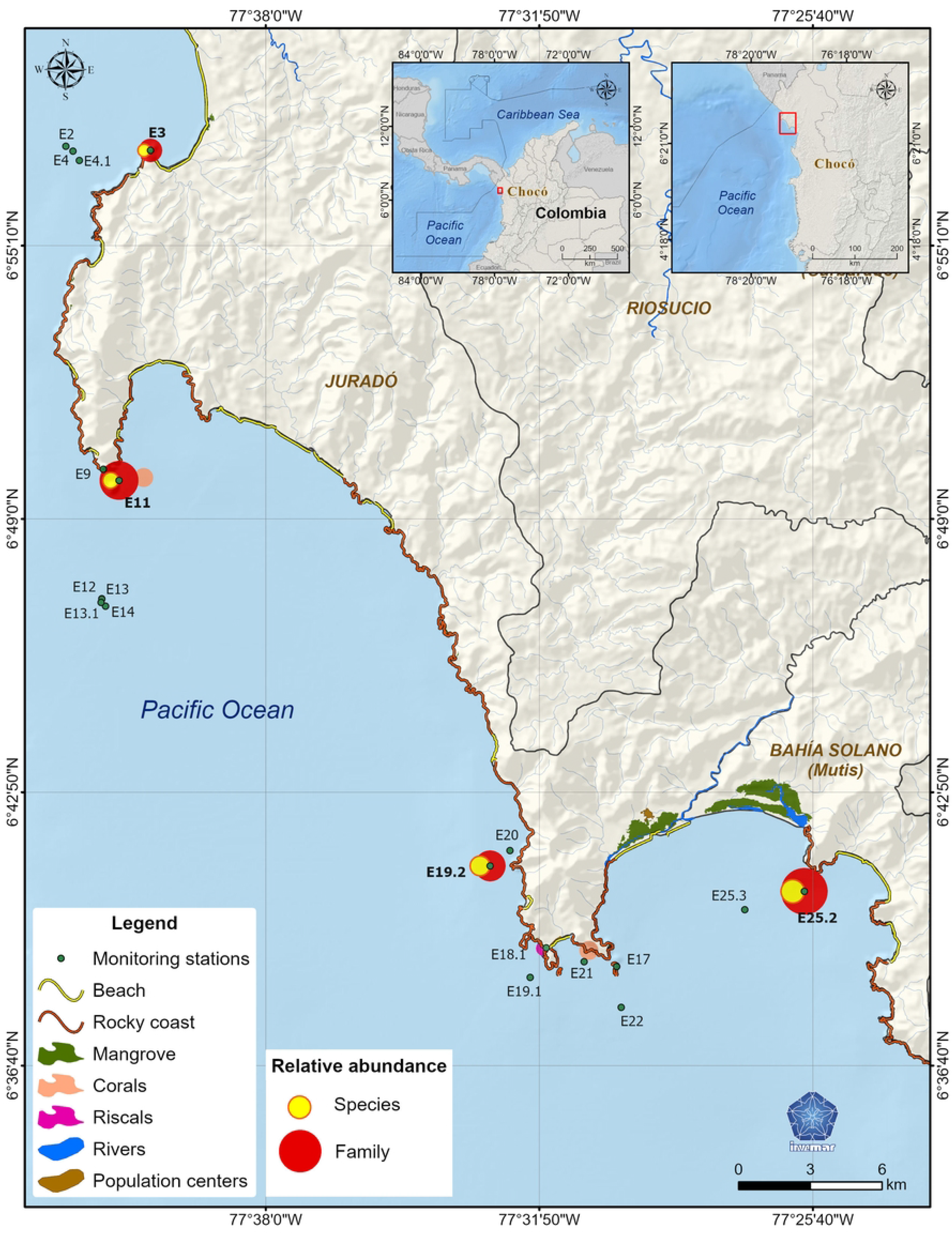
Sampling stations. Spatial distribution of DNA-based sampling and of taxa composition along rocky habitats between Juradó municipality and Cupica Gulf. Map developed by the Information Services Laboratory of INVEMAR (LABSIS).

Marine ecosystems comprise rocky reefs (<60 ha) and small coral assemblages that sustain highly diverse benthic and pelagic communities, serving as critical habitats for reef fishes, invertebrates, and macroalgae [10, 30]. These systems function as biological corridors for seabirds and migratory species, support artisanal fisheries targeting Lutjanidae, Serranidae, and Thunnini [31,28], and act as fish aggregation sites hosting coral and habitat-forming invertebrates [10,20].

### Environmental DNA sampling, metabarcoding and bioinformatics

Aquatic eDNA sampling was conducted along four longitudinal transects adjacent to rocky reef formations (Fig 1; S1 Table). Seawater was collected during high tide in daylight from the bow of a 3 m vessel. At each site, two Athena® peristaltic pumps (Proactive Environmental Products LLC) were deployed on opposite sides of the vessel, each connected to sterile 2 m-long silicone tubing and single-use 0.2 μm filtration capsules (VigiDNA; SPYGEN). Filtration occurred at 1.5 m depth for 30 min at a flow rate of 1 L min⁻¹, yielding two replicates of 30 L per site (60 L total). Sampling was performed from a moving vessel traveling at 2 knots along longitudinal transects parallel to the coastline. Following filtration, residual seawater retained in the capsule was expelled, and eDNA was stabilized by adding 80 mL of CL1 preservation buffer (SPYGEN). Capsules were then stored at ambient temperature until subsequent laboratory processing.

eDNA extraction and PCR amplification were performed on 12 replicates per sample using the primer sets Vert01, Euka03, and Tele01 (Table 1), following standardaized protocols [32,33]. Purified PCR amplicons were pooled in equimolar concentrations to target ∼1,000,000 reads per sample. Three libraries were prepared using Illumina TruSeq and sequenced on an Illumina NextSeq 1000 (paired-end, 2 × 150 bp) following manufacturer protocols. Negative controls (extraction and PCR blanks) were processed and sequenced alongside samples.

**Table 1.**
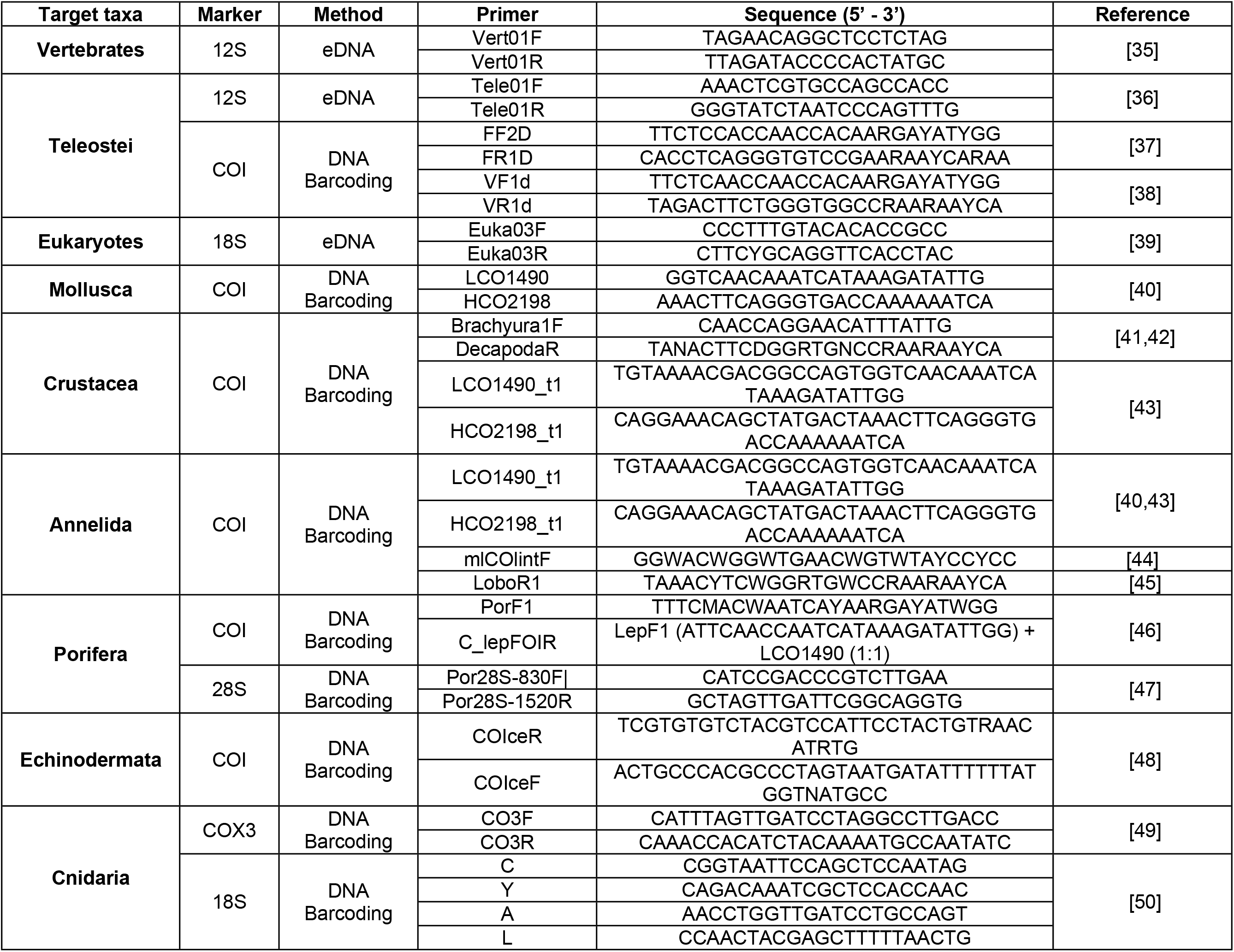
DNA Barcoding and eDNA metabarcoding markers used.

Bioinformatic processing were conducted using OBITools pipeline [34,36] then we performed paired-end merging (alignment score ≥40) and demultiplexing (NGSFILTER). Subsequently, individual datasets were generated with OBISPLIT for downstream processing. Identical sequences were clustered (OBIUNIQ) and sequences were filtered (<20 bp or <10 occurrences removed with OBIG-REP). ASV taxonomic assignments employed ECOTAG against GenBank® release 247 with identity thresholds of 98–100% (species), 96–98% (genus), and 90–96% (family). All other ASVs assigned to higher categories were removed from subsequent analysis. Taxonomic annotations were validated against regional and global reference databases, including OBIS, GBIF, FishBase, and the World Register of Marine Species (WoRMS). Queries were performed using species names and higher taxonomic ranks derived from sequence assignments. Each record was cross-checked for synonymy, distribution range, and habitat information to ensure consistency with documented coastal metazoan biodiversity of the Eastern Tropical Pacific. Terrestrial eukaryotes and non-metazoan detections were also eliminated from the analysis.

### Metazoan sampling and DNA Barcoding

Metazoan specimens were manually collected from 14 rocky subtidal sites through SCUBA diving (S1 Table). The collection comprised 769 individuals representing 11 phyla: Arthropoda (21.42%), Echinodermata (19.66%), Mollusca (17.33%), Chordata (13.6%), Annelida (8.2%), Porifera (6.4%), Cnidaria (5.4%), Brachiopoda (3.5%), Nemertea (2.4%), Platyhelminthes (1.09%), and Bryozoa (1%). Specimens were anesthetized with magnesium chloride solution to preserve structural integrity prior tissue sampling. From each individual, tissue subsamples (1-2 cm) were excised and preserved in molecular-grade ethanol. Voucher specimens were photographed, and associated metadata were recorded before fixation in 70% ethanol or formalin (for fish). To ensure DNA integrity, ethanol in tissue samples was replaced at 6, 12, and 24 hours post-preservation, after which samples were stored at −20 °C until laboratory processing.

Genomic DNA was isolated from all phyla using a modified salt-extraction protocol [51] and the DNeasy Tissue and Blood Kit (QIAGEN), following manufacturer guidelines with minor modifications. DNA quality and quantity were assessed based on concentration and purity ratios (260/280 and 260/230). Samples with concentrations >20 ng/µL, 260/280 ratios of 1.8-2.0, and 260/230 ratios of 2.0-2.2 were retained and stored at −20 °C. Fragments >500 bp were amplified in 30 µL reactions using primers targeting COI, 28S, and 18S genes (Table 1). PCR mixtures contained 1× Taq buffer, 2 mM MgCl₂, 0.2-0.4 mM dNTPs, 0.1 µM of each primer, 0.1 U RedTaq polymerase, and 1-3 µL of template DNA. Thermal cycling consisted of an initial denaturation at 94 °C for 2 min, followed by 35 cycles of 94 °C for 30 s, 52-55 °C for 40 s, and 72 °C for 1 min, with a final extension at 72 °C for 5-10 min.

Positive PCR products were sequenced using Sanger technology (forward and reverse reads). Raw sequences were quality-checked and assembled into consensus sequences using Geneious Prime (Dotmatics). Taxonomic assignments for each OTU were validated by expert taxonomists and through NCBI BLASTn and BOLD sequence matching algorithms by applying similarity thresholds of >99% for species, 90-99% for genus, and 85-90% for family-level identifications.

Specimen and environmental DNA (eDNA) sampling conducted by INVEMAR did not require a collection permit, in accordance with the Colombian regulatory framework. Specifically, as established in Paragraph 1, Article 2.2.2.8.1.2., Section 1 (Permits), Chapter 8 (Scientific Research) of Decree 1076 of 2015, entities affiliated with or linked to the Ministry of Environment and Sustainable Development, including INVEMAR, are exempt from the requirement to obtain permits for the collection of wild specimens. Given INVEMAR’s status as a linked entity to the Ministry (Article 1.2.2.1., Title 2, Decree 1076 of 2015), all specimen and eDNA collections carried out under this study were legally performed without the need for a collection permit.

### Data analysis

Metazoan diversity was assessed at phylum, order, family, and ASV (Amplicon sequence variant) levels. Due to incomplete reference libraries for the Eastern Tropical Pacific, ∼25% of the taxa could be identified to species level with the combination of specimen morphological taxonomy and DNA approaches, therefore, most sequences were resolved to family level to reduce misclassification risk. Higher taxonomic ranks ensured robust site comparisons, while ASV-level analyses, independent of taxonomic annotation, explored fine-scale diversity patterns. Data from eDNA sequences were normalized by converting read counts per phylum per site into relative frequencies.

All statistical analyses were conducted in R version 4.5.1. Visualizations, including bar plots and Venn diagrams, were generated using the *ggplot2* and *VennDiagram* (v1.6.2) packages. Sequence reads from eDNA metabarcoding across sampling sites and DNA barcoding were pooled at the species level for analysis. To determine the effect of sampling efforts to overall detected richness, ASV-level rarefaction curves were created for each sampling site using “*specaccum*” function in the *Vegan* package, data from specimen collection was not pooled against eDNA due to larger sampling effort. PERMANOVA (10,000 permutations) were performed to test the effect of sampling site on ASVs taxonomic composition. Mann-Whitney-Wilcoxon test was performed to compare or differences in taxonomic resoluti between methods.

Alpha diversity indices (Shannon (H′) and Simpson indices) were calculated with the *BiodiversityR* package. Shannon index was selected because it accounts for both species richness and evenness, providing a measure of taxonomic complexity, while the Simpson index emphasized dominant taxa, offering insights into species dominance patterns. Differences in ASV richness between sampling sites and methods were tested using the Kruskal–Wallis test (95% CI).

Beta diversity was quantified using Bray-Curtis dissimilarity with the beta.div.comp function in the *vegan* package and the incidence of Jaccard coefficient was chosen because it is a presence-absence-based metric, making it suitable for eDNA data where detection probabilities vary and abundance estimates may be biased. This approach emphasized compositional differences rather than read count variability, providing a robust measure of community turnover among sites.

### Ethical statement

The authors declare that they all agree with this publication and made significant contributions; that there is no conflict of interest of any kind; and that we followed all pertinent ethical and legal procedures and requirements.

## Results

### eDNA metabarcoding

Amplicon sequencing targeting the Vert01, Tele01, and Euka03 loci generated a total of 7,572,247 high-quality reads following bioinformatic filtering (S2-4 Tables). These reads were assigned to 1,444 molecular operational taxonomic units (MOTUs) after the exclusion of non-marine taxa (mean = 4,580 reads per sample; SD = 487.37). Subsequent data curation involved the removal of low-abundance features (<50 reads) to mitigate the influence of spurious detections and minimize the overinterpretation of stochastic amplification artefacts. This filtering step retained 3,357,922 reads and 1,122 MOTUs for downstream analyses.

Across primer sets results revealed differential performance in read yield and taxonomic resolution. Euka03 generated 236 MOTUs, of which 211 were assigned to the order level or higher taxonomic ranks, accounting for 710,349 reads (mean = 3,035 reads per MOTU). The Tele01 marker yielded 196 MOTUs, with 72 resolved to species level, representing 2,549,892 reads (mean = 13,009 reads per MOTU). In contrast, the Vert01 marker produced the highest diversity, with 690 MOTUs, including 88 assignments at species level, and a total of 4,312,006 reads (mean = 6,249 reads per MOTU).

Taxonomic assignments spanned multiple hierarchical levels, including species, genus, family, and higher taxonomic categories (e.g., tribe and infraorder). However, species-level resolution remained constrained due to the incomplete representation of 12S and 18S V7 reference databases (S5 Table), resulting in the majority of MOTUs being assigned to higher taxonomic ranks. Overall, the dataset comprised classifications across 13 phyla, 25 classes, 83 orders, 83 families, 118 genera, and 106 taxa identified to the species level.

### DNA Barcode database

A total of 1,460 tissue subsamples were processed from 769 individual specimens. Of these, 280 samples (19%) failed to meet the minimum quality and quantity thresholds for DNA extraction and amplification and were excluded from downstream analyses. The barcoding effort yielded 871 high-quality consensus sequences, of which 91.6% originated from the COI region (>600 bp), resulting in 414 validated barcodes from 178 OTUs (belonging to 76 species and 64 families). The remaining 8.3% corresponded to sequences from the 18S (24 sequences) and 28S (49 sequences) regions; however, these did not produce complete barcodes. Because eDNA metabarcoding was not performed using 18S or 28S amplicons, barcodes generated from these loci were not incorporated into reference databases for direct comparison with eDNA datasets. Nevertheless, all sequences have been deposited in the BOLD Systems database under the CCBIO (Colombia BIO) project container to ensure public accessibility. A detailed summary of taxa successfully barcoded using COI fragments is provided in S6 Table.

### Taxonomic composition and richness

The combination of visual censuses, specimen collection with morphological identification, DNA barcoding based on COI sequences, and eDNA metabarcoding yielded unique 1,300 taxonomic units, asymptote in the rarefaction and extrapolation curves (95% CI) suggested that the sequencing depth and sampling effort captured most of the community diversity (Fig 2).

**Fig 2.**
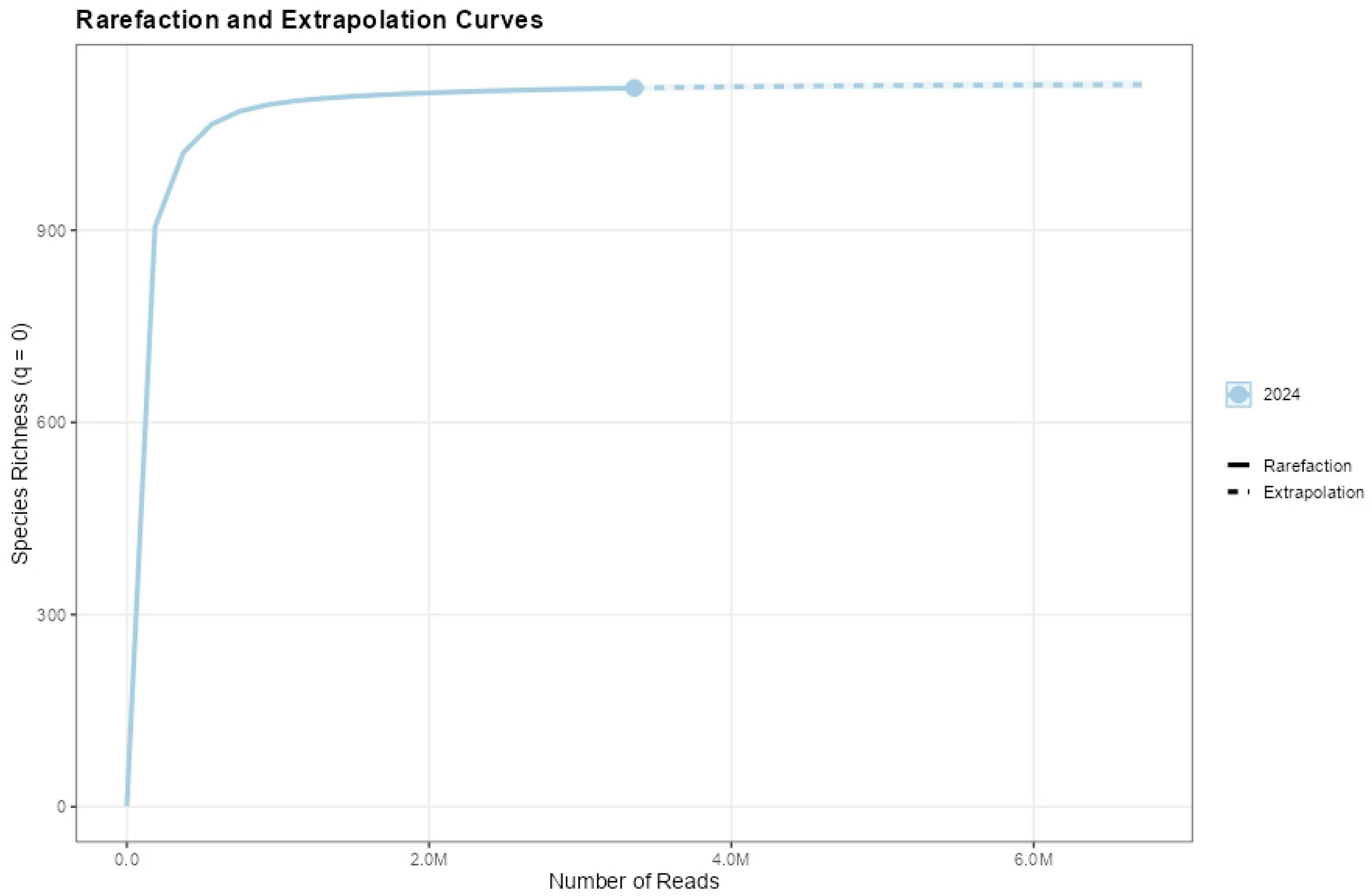
Rarefaction and extrapolation curves. Asymptote obtained indicating sufficient sampling and sequencing depth.

The taxonomic resolution of eDNA per sample (mean = 58.2) was significantly higher than through DNA barcoding (mean= 17; Mann-Whitney-Wilcoxon test W = 85.4, p< 0.05). Gamma diversity was higher for eDNA despite greater sampling effort for specimen collections. Chordata (Teleostei) was the most abundant phylum in both datasets; however, relative taxon abundance differed significantly between methods (S2-6 Tables, S8 Fig).

Primer performance varied across sites, Vert01 exhibited the highest taxonomic resolution at sites E11 (310 ASVs) and E25 (397 ASVs), outperforming Tele01 (137 ASVs; and 130 ASVs respectively) (Fig 3). Site E19 showed the greatest invertebrate diversity with Euka03, while overall diversity was lowest at E3 across all primers (S7 Table). We found that richness varied across sampling sites and was higher in E25 (p<0.001) and the lowest in E3 (p<0.001). Taxonomic orders detected via eDNA were consistent with those observed through specimen collections (Fig 3). Fish identification made with eDNA metabarcoding and DNA barcoding included, solitary (56%), social (20%), commercially important species (15%) and rare (9%) species.

**Fig 3.**
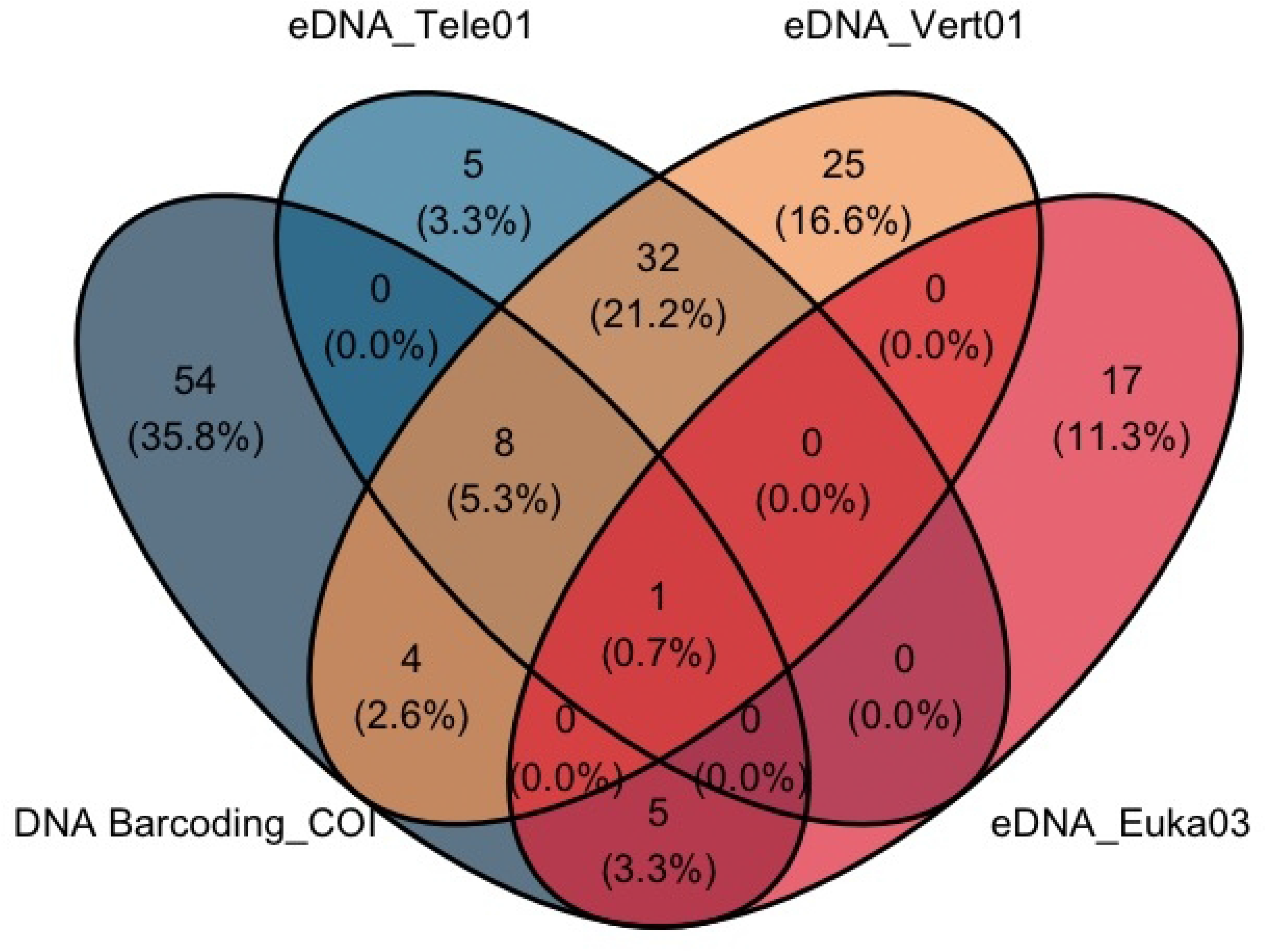
Taxa detections across DNA primers. Venn diagram showing the number of overlapping ASVs between DNA Barcoding_COI, and eDNA (Primers Tele01, Euka03 and Vert01).

Phylum-level taxonomic composition obtained with eDNA metacarcoding was strongly dominated by Chordata, which accounted for 83% of total relative abundance (mean = 72% ± 40), followed by Arthropoda (∼12%; mean = 21% ± 32). Mollusca (mean = 3.6% ± 7.2) and Cnidaria (mean = 1.6% ± 3.5) ranked as the third and fourth most abundant phyla, contributing 2.1% and 0.83%, respectively (Fig 4).

**Fig 4.**
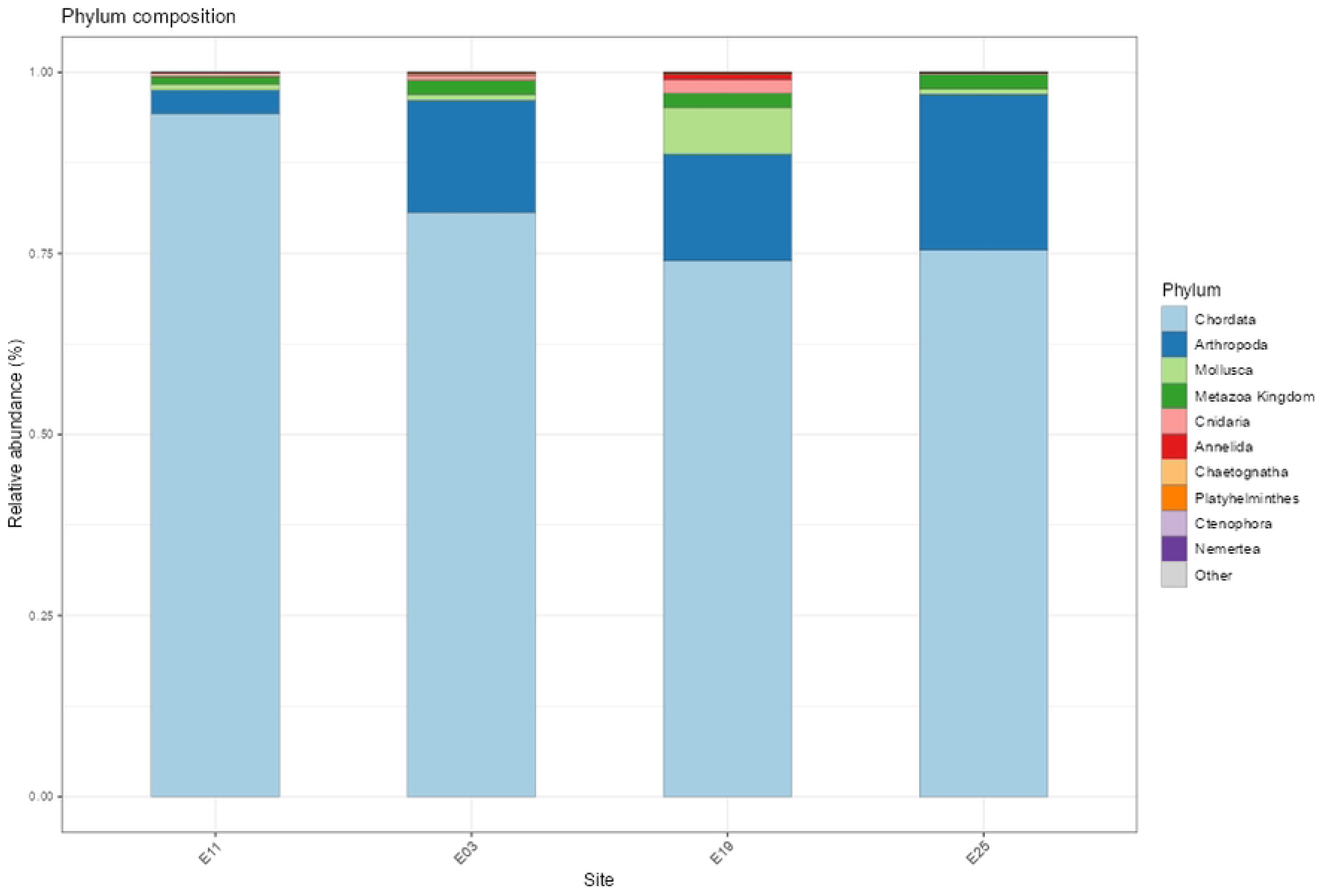
Taxonomic composition. Phylum level composition obtained with eDNA-based approaches.

Spatial variation was evident among sampling sites. At site E11, Chordata reached its highest relative abundance (95.3%), while Arthropoda showed its lowest contribution (3.2%). In contrast, Arthropoda peaked at site E03 (25.7%), where low-abundance phyla such as Chaetognatha (0.27%) and Brachiopoda (0.01%) were also detected. Maximum relative abundances for Mollusca (7.2%) and Annelida (0.84%) were observed at site E19. Site E25 exhibited the highest relative abundance of Nemertea (0.03%), the second-highest representation of Chordata, and among the lowest contributions of several low-abundance phyla, including Annelida (0.06%), Chaetognatha (0.01%), and Platyhelminthes (0.01%).

Notably, Cnidaria displayed the greatest variability across sites, ranging from 0.18% (E25) to 3.3% (E11). Overall, phylum-level patterns indicate a vertebrate-dominated assemblage, while variability in Mollusca, Arthropoda, and other low-abundance phyla suggests site-specific community structure, likely influenced by differences in microhabitat composition (Fig 4).

Class-level composition was dominated by Actinopteri (79%; mean = 63% ± 46) and Hexanauplia (12%; mean = 21% ± 32), which together accounted for the majority of reads and reflected the prevalence of Chordata and Arthropoda. Minor contributions from Appendicularia (3.6%; mean = 5.7% ± 8.8) and Gastropoda (1.9%; mean = 3.5% ± 7.1) resulted in these four classes comprising 97.7% of total abundance, with moderate spatial variability observed across sites (Fig 5).

**Fig 5.**
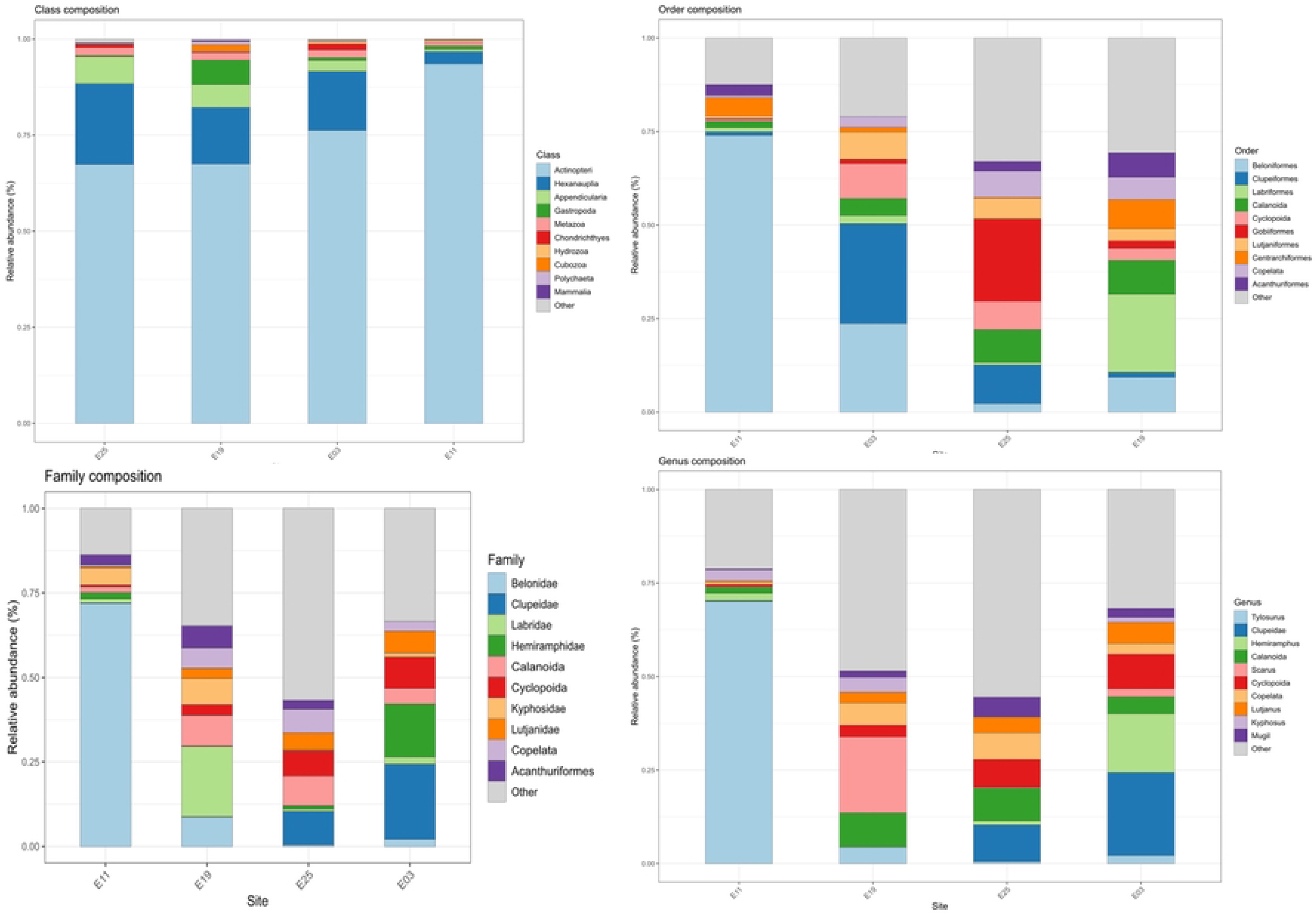
Comparison of the taxonomic composition. Class, Order, family and genus level composition obtained with eDNA-based approaches.

At finer taxonomic resolution, order-level composition exhibited greater site-specific differentiation. Beloniformes was the dominant order (32%; mean = 13.7% ± 22.8), followed by Clupeiformes (10.7%; mean = 9.3% ± 11.8) and Labriformes (5.8%; mean = 3.6% ± 11.0). Among arthropods, only Calanoida (5.5%; mean = 9.7% ± 14.0) and Cyclopoida (5.1%; mean = 8.0% ± 14.8) contributed substantially, with Calanoida showing relatively consistent abundance across sites (Fig 5). These patterns indicate that ecological variability becomes more evident at lower taxonomic levels.

Family-level analysis revealed strong dominance by teleost fishes, with the ten most abundant families accounting for 78.3% of total reads. Notably, Belonidae (30%; mean = 14% ± 25) and Clupeidae (10.9%; mean = 8.3% ± 10.8) were the most abundant and variable families (Fig 5, S7 Table). At the genus level, Tylosurus was the most dominant taxon (39.2%; mean = 14% ± 27), followed by Hemiramphus (9.5%) and Scarus (8.9%). Marked spatial heterogeneity was observed, with strong dominance of Tylosurus at site E11 (∼80%), Hemiramphus at E03 (∼39%), and Scarus at E19, whereas E25 exhibited a more even assemblage (Fig 5). The high standard deviations, particularly for dominant taxa, further highlight substantial spatial variability in community structure, suggesting habitat-driven differences among sites.

Mean observed MOTU richness across samples was 90.58 (±52.24), ranging from 32 (E03_83_t) to 244 (E25_86_v). Mean Shannon diversity and Inverse Simpson indices were 2.95 (±0.85) and 15.49 (±12.51), respectively. Among sites, E25 exhibited the highest mean richness (111 MOTUs) and Shannon diversity (3.26), indicating comparatively higher diversity, whereas E03 showed the lowest mean richness (64.67 MOTUs). Site E11 displayed the greatest variability in diversity metrics (Observed ±53.72; Inverse Simpson ±19.89), with some samples characterized by low evenness (minimum Inverse Simpson = 1.46; Shannon = 1.04), suggesting dominance by few taxa. In contrast, E19 exhibited relatively higher and more consistent diversity. Despite this variability, Kruskal–Wallis tests indicated no significant differences among sites for observed richness, Shannon diversity, or Inverse Simpson indices (all p > 0.05), suggesting comparable alpha diversity across locations. Overall, the study area exhibited substantial heterogeneity in community structure, with E25 and E11 supporting comparatively richer but more variable assemblages, whereas E03 and E19 showed lower variability and more homogeneous community composition (Fig 6).

**Fig 6.**
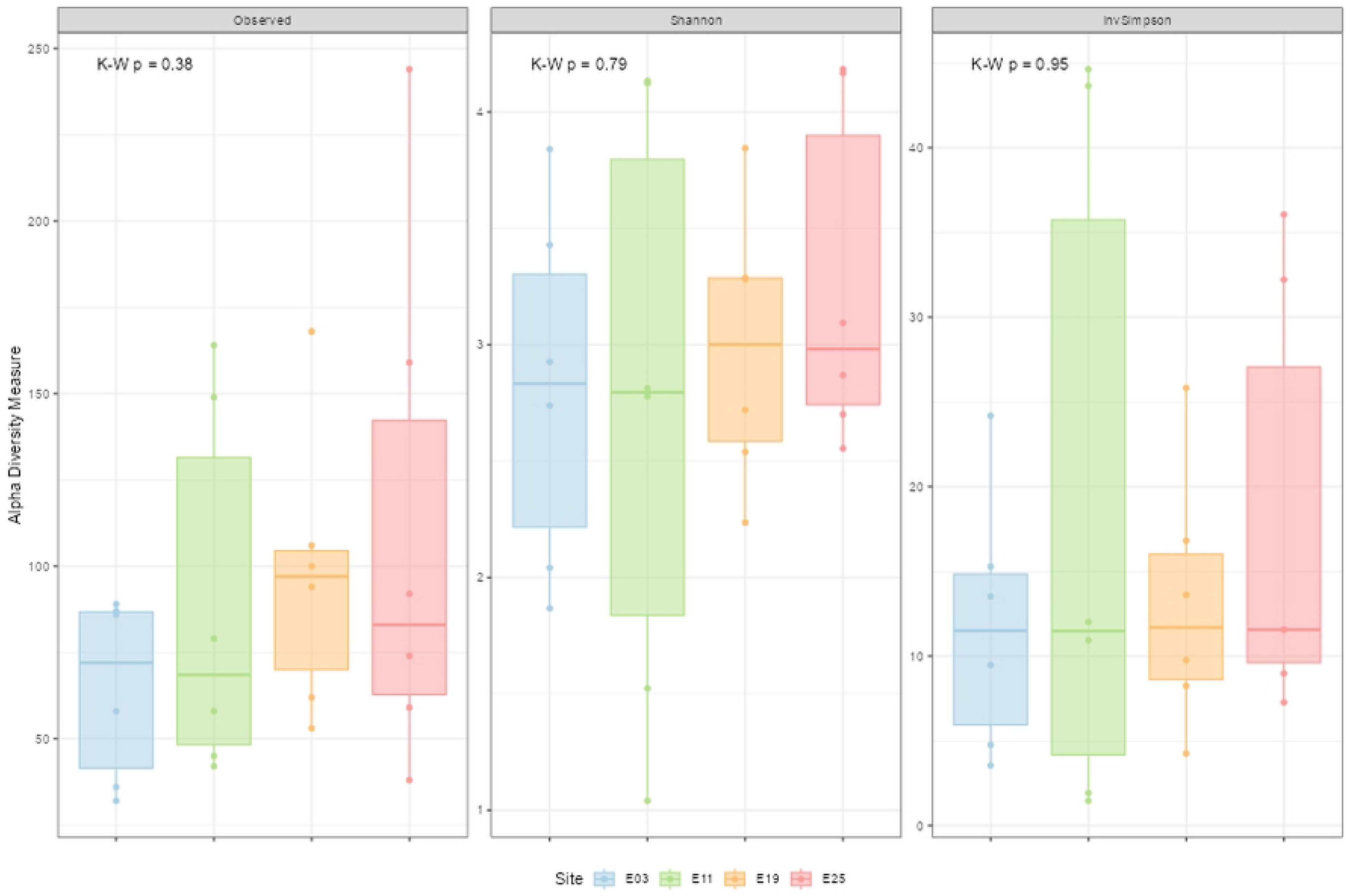
Observed richness, Shannon, and Inverse Simpson indices across sites (mean ± SD). No significant differences among sites (Kruskal–Wallis, p > 0.05), though variability was highest at E11 and diversity peaked at E25.

Beta diversity analyses based on abundance (Bray–Curtis; PERMANOVA, p = 0.26; dispersion test, p = 0.95) and incidence (Jaccard; PERMANOVA, p = 0.47; dispersion test, p = 0.92) revealed no statistically significant differences in community structure among sampling sites. Variance explained by site was low, accounting for 14% (Bray– Curtis) and 12% (Jaccard), with the majority of variation attributable to within-site heterogeneity. Partitioning of total beta diversity (βₛₒᵣ = 0.95) indicated that community dissimilarity was overwhelmingly driven by species turnover (βₛᵢₘ = 0.93; 97.7%), with a minor contribution from nestedness (βₛₙₑ = 0.02; 2.3%). These results suggest that the sampled locations collectively represent a single, species-rich and highly heterogeneous community, structured primarily by species replacement rather than nested subsets of taxa.

## Discussion

### Subtidal metazoan diversity

Subtidal habitats are permanently submerged environments characterized by relatively stable physical conditions, low turbidity, and high light penetration, which support the proliferation of macroalgae and other primary producers [52,53]. Hard substrates provide essential attachment surfaces for sessile organisms, while structural complexity offers refugia for mobile invertebrates and fish, contributing to higher biodiversity [54]. These ecosystems sustain intricate trophic networks comprising herbivores, detritivores, predators and filter feeders playing a pivotal role in nutrient cycling and benthic-pelagic coupling [55]. Hydrodynamic regimes strongly influence species distribution, recruitment, and community structure, while connectivity with adjacent ecosystems enhances their ecological significance as biodiversity reservoirs and functional corridors within coastal landscapes [56].

This study integrated eDNA metabarcoding and specimen-based DNA barcoding to enhance metazoan inventories associated with rocky subtidal habitats in the northern Colombian Pacific. Despite the limited representation of marine taxa in genetic reference databases for the Eastern Tropical Pacific (particularly in the northernmost Colombian region) eDNA metabarcoding consistently recovered a broader taxonomic record compared to traditional specimen collections, even with greater sampling effort for the latter. These findings highlight the capacity of eDNA approaches to detect cryptic and low-abundance taxa, thereby improving biodiversity assessments in data-poor regions [57].

We found that the combination of different approaches led to an improved metazoan inventory encompassing taxa from small invertebrates to rare and cryptic fish species as well as marine mammals. Species detected were primarily associated with rocky reef subtidal habitats or exhibited demersal life stages. Among all fish species identified, 20 taxa accounted for the highest read abundance across sampling sites and were consistently detected by both eDNA metabarcoding and DNA barcoding approaches. These included demersal species such as *Synodus evermanni* and *Erotelis armiger*; pelagic-oceanic taxa (*Platybelone pterura*, *Tylosurus pacificus*); reef-associated species (*Tylosurus melanotus, Entomacrodus chiostictus, Hypsoblennius brevipinnis, Decapterus macrosoma, Selar crumenophthalmus, Trachinotus rhodopus, Lutjanus guttatus, Scarus rubroviolaceus, Acanthemblemaria balanorum, Bathygobius ramosus, Gobiesox adustus, Protemblemaria bicirrus*); pelagic-neritic taxa (*Caranx caballus, Opisthonema libertate*); and rocky reef specialists (*Dialommus macrocephalus, Elacatinus puncticulatus*).

Rock-associated invertebrates were also detected with both approaches, the greatest read abundance were assigned to mollusks, such as chitons (*Callistochiton pulchrior, Ischnochiton dispar, Chiton stokesii*), gastropods (*Bostrycapulus aculeatus, Crucibulum scutellatum*), bivalves (*Pinctada mazatlanica*), and opisthobranchs (*Chromolaichma dalli*). Echinoderms such as *Diadema mexicanum* (sea urchin), *Holothuria (Halodeima) inornata* (sea cucumber), and *Ophiocomella alexandri* (brittle star), and to decapod crustaceans (*Eriphia squamata, Eupilumnus xantusii*).

The metazoan diversity documented in this study represents 48.5% of all taxa previously reported for the Colombian Pacific. The Jurado–Cupica stretch, corresponds to 15% of the national Pacific coast in which we recorded 83 orders, 125 families, 106 genera, and 175 species. These findings constitute the first DNA-based faunal inventory for the northern Colombian Pacific and the most comprehensive dataset generated to date. The limited availability of biodiversity records for the Eastern Tropical Pacific has been widely documented in the literature [2,58]. Updated taxonomic checklists and inventories for rocky subtidal fauna are particularly scarce [15], with most existing efforts concentrated on beach meiofauna and mangrove-associated invertebrates and fishes in the mid-southern coastal region 6,19,20,21]. This lack of comprehensive inventories for subtidal ecosystems underscores a critical knowledge gap that hampers effective biodiversity assessments and conservation planning in the region.

Expanding species inventories in this region is critical for improving baseline biodiversity knowledge, particularly in understudied ecosystems such as rocky subtidal habitats. Robust inventories are essential for informing conservation strategies, guiding marine spatial planning, and assessing ecosystem resilience under increasing anthropogenic pressures and climate change. The integration of molecular tools, such as eDNA metabarcoding and DNA barcoding, provides an effective approach to overcome taxonomic gaps and detect cryptic or rare species, thereby strengthening biodiversity monitoring frameworks in data-poor regions.

### Completeness of eDNA-based biodiversity inventories and sample collections

eDNA metabarcoding provided extensive taxonomic coverage, detecting species across pelagic, neritic, and demersal zones, thereby capturing a broad representation of ecological niches. In contrast, DNA barcoding offered higher taxonomic resolution at the species level, particularly for sessile and low-frequency taxa such as rocky-adhered mollusks and echinoderms. This complementarity highlights the strengths of integrating both approaches for comprehensive biodiversity assessments in structurally complex subtidal ecosystems.

However, species detection through eDNA metabarcoding is strongly influenced by local oceanographic conditions, as the transport and persistence of DNA fragments vary according to hydrodynamic processes, water column stratification, and ecosystem dynamics [59,60]). These factors affect the spatial and temporal distribution of eDNA, potentially introducing biases in species detection. Furthermore, taxonomic resolution is contingent upon primer specificity, which determines amplification success across different taxa [61]. Consequently, both environmental variability and primer design play critical roles in shaping the completeness and accuracy of eDNA-based biodiversity assessments [62,63].

Despite its utility, DNA barcoding is constrained by several limitations. Its accuracy relies heavily on the completeness of reference databases, which remain insufficient for many taxa in the Eastern Tropical Pacific [12]. This gap often results in higher-level taxonomic assignments rather than species-level identifications. Furthermore, specimen-based methods are labor-intensive, require physical collection, and are biased toward larger or more conspicuous organisms, potentially underrepresenting cryptic or rare species [33,64]. These limitations underscore the importance of combining DNA barcoding with eDNA metabarcoding to overcome taxonomic gaps and improve inventory completeness.

The ability of eDNA metabarcoding to detect a wide range of taxa, including elusive and low-abundance species, demonstrates its potential as a powerful tool for biodiversity monitoring in data-poor regions. However, the limited representation of regional taxa in existing reference databases has likely contributed to the observed discrepancies between DNA barcoding and eDNA metabarcoding results, particularly at finer taxonomic levels. In response to this limitation, we generated a novel DNA barcode dataset based on COI sequences derived from rigorously identified specimens collected during the study. These specimens were identified using standard taxonomic procedures by expert taxonomists and have been deposited in recognized Natural History Museum collections to ensure traceability and support future research efforts.

While the generation of additional reference sequences is expected to improve congruence between molecular approaches, such efforts require sustained sampling and sequencing initiatives beyond the scope of the present study. Nonetheless, the dataset produced here constitutes a significant step toward strengthening regional reference libraries, thereby contributing to the progressive improvement of molecular biodiversity assessments in the Colombian Pacific

Continued efforts to expand reference libraries are essential to maximize species-level resolution and fully leverage the advantages of both molecular approaches that led to the creation of the first mitocondrial database of marine fauna in the region.

Differences in species detection between eDNA metabarcoding and specimen-based collection are also influenced by taxon-specific variability in detectability across these approaches. Certain taxa were exclusively identified through physical specimen collection, whereas others were detected only sporadically or at low frequency using eDNA. These discrepancies may reflect differences in organismal abundance, DNA shedding rates, and habitat association, as well as methodological factors such as primer specificity and amplification efficiency. Understanding these detection biases is essential for interpreting biodiversity patterns and optimizing integrated monitoring strategies.

These species reflect the structural complexity and ecological diversity of rocky reef habitats, encompassing key functional groups such as grazers, filter feeders, and benthic predators and life-history strategies. Their detection by both molecular approaches underscores the complementarity of eDNA and specimen-based barcoding for biodiversity assessments in structurally complex marine ecosystems.

The molecular baseline generated in this study provides a critical reference framework for detecting and quantifying biodiversity change in a region highly vulnerable to climate-driven impacts. The high taxonomic richness, coupled with substantial spatial turnover (βₛᵢₘ = 97.7%), indicates that local assemblages are strongly structured by environmental heterogeneity, suggesting potential sensitivity to shifts in oceanographic conditions associated with global warming. Under scenarios of increasing sea surface temperature, altered hydrodynamics, and ecosystem simplification, such turnover-driven communities may undergo rapid compositional reorganization before detectable losses in alpha diversity occur.

By integrating eDNA metabarcoding with specimen-based DNA barcoding, this study also contributes to addressing one of the primary limitations in tropical biodiversity research: the incomplete representation of taxa in reference databases. The generation of region-specific molecular resources enhances taxonomic resolution and enables more accurate long-term monitoring, which is essential for detecting early warning signals of biodiversity erosion.

At a broader scale, these results support the implementation of cost-efficient, high-resolution molecular monitoring frameworks in data-poor regions, strengthening capacity for adaptive management and conservation planning. In the context of accelerated biodiversity loss, such integrative approaches are fundamental to bridging knowledge gaps, improving detection of cryptic and declining taxa, and informing policy decisions aimed at mitigating the ecological and socio-economic consequences of marine biodiversity decline in the Eastern Tropical Pacific.

## Conclusions

This study provides the first DNA-based faunal inventory for rocky subtidal habitats in the northern Colombian Pacific and represents the most comprehensive dataset available for the Jurado–Cupica region. By integrating eDNA metabarcoding with specimen-based DNA barcoding, we substantially expanded metazoan inventories, capturing taxa ranging from small, substrate-associated invertebrates to rare and cryptic fishes and marine mammals, while documenting pronounced spatial variability in taxonomic composition. eDNA metabarcoding consistently achieved broader taxonomic coverage and enhanced detection of low-abundance and elusive species, whereas DNA barcoding provided higher taxonomic resolution, particularly for sessile and infrequent taxa. These results underscore the complementarity of both approaches in structurally complex tropical ecosystems.

The resulting inventory accounts for 48.5% of all taxa previously reported for the Colombian Pacific and establishes a robust molecular baseline for biodiversity monitoring, marine spatial planning, and resilience assessments under increasing anthropogenic and climatic pressures. Nonetheless, methodological and infrastructural limitations remain. eDNA-based detections are influenced by hydrodynamic processes and primer biases, while DNA barcoding resolution is constrained by incomplete reference libraries. Addressing these challenges through expanded regional genomic resources, standardized methodologies, and sustained monitoring efforts will be essential to improve taxonomic resolution and detection accuracy. The use of multiple genetic markers combined with specimen-based approaches proved critical to maximizing biodiversity detection and should be considered a standard practice in future surveys.

Beyond its regional implications, this study contributes a scalable framework for biodiversity assessment in data-limited tropical marine systems. The high species turnover observed across sites highlights the ecological sensitivity of these assemblages to environmental change, suggesting that climate-driven shifts may result in rapid compositional reorganization even in the absence of immediate declines in overall diversity. In the context of accelerated global biodiversity loss, the generation of standardized, high-resolution baseline datasets such as this is essential for detecting early ecological changes, identifying vulnerable taxa, and informing adaptive conservation strategies. By strengthening molecular reference libraries and demonstrating the effectiveness of integrative approaches, this work supports the development of cost-efficient, reproducible monitoring systems capable of informing policy, mitigating biodiversity loss, and enhancing ecosystem resilience in the Eastern Tropical Pacific and other underexplored regions worldwide.

## Supporting information

**S1 Table. Details of sampling sites.** Sampling sites for all complementary DNA-based approaches.

**S2 Table. Taxa composition and distribution for vertebrates.** ASV Taxonomic resolution using VERT01 primer

**S3 Table. Taxa composition and distribution for fish.** ASV Taxonomic resolution using TELE01 primer

**S4 Table**. **Taxa composition and distribution for eukaryonts.** ASV Taxonomic resolution using EUKA03 primer

**S5 Table. Species-level composition and distributions.** Taxonomic classification for all species-level detections

**S6 Table. DNA Barcodes created in BOLDSystems.** Taxonomic resolution from tissue samples using COI primer

**S7 Table. Famly-Level composittion and distribution.** Number of Family-level seq reads per sampling site (after bioinformatic filters)

**S8 Fig. Barplot of comparisons between taxa composition.** Taxonomic composition at the phylum level for eDNA and DNA barcoding approaches.

## Data availability

All biodiversity metadata including morphological metazoa data from the Expedition BIO Jurado-Cupica is published as a datset at https://www.gbif.org/dataset/6e5801dd-296a-4318-95ce-64ea11e7efc8 and https://obis.org/dataset/36ebf033-10b8-42a8-ad21-38b1b78d1867. eDNA-derived data is published in OBIS through a restricted mode and will be made publicly available upon acceptance for publication. On the other hand, Mitochondrial barcodes are uploaded on BOLDSystems database inside the CCBIO container (Colombia BIO), under the following projects: Colombia-BIO-INVEMAR-Peces-Jurado (CBIPJ) (Fish), Colombia-BIO-INVEMAR-Crustacea-Jurado (CBICJ) (Crustaceans), Colombia-BIO-INVEMAR-Anelidos-Jurado (CBIAJ) (Annelids), Colombia-BIO-INVEMAR-Equinodermos-Jurado (CBIEJ) (Echinoderms) Colombia-BIO-INVEMAR-Moluscos-Jurado (CBIMJ) (Mollusks) and Colombia-BIO-INVEMAR-Reptilia-Jurado (CBIRJ) (Reptiles) (DNA sequences will be made public upon manuscript acceptance for publication).

## Acknowledgments

Special thanks to the Los Delfines Community Council Collective (CCG Los Delfines) and the Cupica Community Council (CC de Cupica) for their participation in the the expedition and allowing us to sample in their territory. Also, to our partner entities, the Pacific Environmental Research Institute (IIAP), Universidad del Valle, Universidad Tecnológica del Chocó, the Marine Natural History Museum of Colombia-MAKURIWA and BIOINNOVA (National Center for Science, Technology, and Innovation for the Sustainable Productive Development of Biodiversity) for their support on the delivery of the project. To the Marine and Coastal Research Institute “José Benito Vives de Andreis”-INVEMAR for their administrative and logistical support specially to the project coordinators David Alonso Carvajal, Edgar Fernando Dorado and Martha Vides Casado. We thank interns, students, researchers, professionals, and technicians involved in the project, for supporting the specimen collections and eDNA sampling. To the Information Services Laboratory (LABSIS, GEZ) for preparing the map and metadata. Contribution No. xx from the Marine and Coastal Research Institute (Number provided upon manuscript acceptamce for publication).

## Author Contributions

**Conceptualization:** Vanessa Yepes Narváez

**Data curation:** Vanessa Yepes Narváez, Alejandro Rodríguez-Sánchez, Mayra Atencia-Galindo.

**Formal analysis:** Vanessa Yepes Narváez, Alejandro Rodríguez-Sánchez

**Visualization:** Alejandro Rodríguez-Sánchez, Mayra Atencia-Galindo.

**Writing – Original draft:** Vanessa Yepes-Narváez

**Writing – review & editing:** Vanessa Yepes-Narváez, Alejandro Rodríguez-Sánchez, Mayra Atencia-Galindo.

